# An analysis of laboratory variability and thresholds for human *in vitro* ADME/PK methods

**DOI:** 10.1101/2022.09.27.509731

**Authors:** Urban Fagerholm

## Abstract

**Introduction:** Various *in vitro* methods are used to measure absorption, distribution, metabolism and excretion/pharmacokinetics (ADME/PK) of candidate drugs and predict and decide whether properties are clinically adequate.

**Methods:** Objectives were to evaluate variability within and between laboratories for commonly used human *in vitro* ADME/PK methods and to explore whether reliable thresholds may be defined. The literature was searched for *in vitro* data for intrinsic metabolic clearance (hepatocyte CL_int_), apparent intestinal permeability (Caco-2 P_app_), efflux ratio (Caco-2 ER), solubility (S) and BCS-class, and corresponding clinical estimates. *In vitro* ADME/PK data for three example drugs (atenolol, diclofenac and gemfibrozil) were used to predict human *in vivo* ADME/PK and investigate whether these would pass a compound selection process.

**Results and Conclusions:** Interlaboratory variability is considerable, especially for f_u_, S, ER and BCS-classification, and on average about twice as high as intralaboratory variability. Approximate mean interlaboratory variability for CL_int_, P_app_, ER and f_u_ (3- to 3.5-fold) appears to be about 2- to 3-fold higher than corresponding interlaboratory variability. Mean and maximum interlaboratory range for CL_int_, P_app_, ER, f_u_ and S are approximately 5- to 100-fold and 50- to 4500-fold, respectively, with second largest range for f_u_ and largest range for S. For one drug, laboratories produced almost 1000-fold different CL_int_ • f_u_-values. It appears difficult/impossible to set clear clinically useful thresholds, especially for CL_int_, ER and S. Poor *in vitro-in vivo* consistency for S and BCS-classification and large portions of compounds out of reach for Caco-2 and conventional hepatocyte assays are evident. Predictions for reference compounds are consistent with inadequate *in vivo* ADME/PK. Ways to improve predictions and compound selection are suggested.

## Introduction

Various *in vitro* methods are used to measure and screen absorption, distribution, metabolism and excretion/pharmacokinetic (ADME/PK) properties of candidate drugs, for example, human hepatocytes for intrinsic hepatic metabolic clearance (CL_int_), Caco-2 cells for apparent intestinal permeability (P_app_) and basolateral to apical/apical to basolateral P_app_-efflux ratio (ER), and water and buffers for solubility (S). It is assumed, and also shown, that these to some extent resemble the *in vivo* situation, and are therefore used to predict corresponding human clinical estimates, exposure profiles and doses.

Compound selection and decision making (stop/go) during the drug discovery phase may be based on rank ordering of important basic ADME/PK-properties (for example, choosing the most permeable and metabolically stable in a series of compounds) or integrated ADME/PK-profiles (for example, selecting the compound with predicted most adequate oral bioavailability (F) and half-life (t_½_) and multiple elimination routes), and/or certain thresholds/cut-off values and classes/categories for ADME/PK.

There is a defined permeability/uptake threshold for the Biopharmaceutics Classification System (BCS) - high permeability when fraction absorbed (f_a_) based on permeation is or is predicted to be ≥90 %, otherwise, low permeability. The BCS also has a cut-off limit for S - high S when the highest dose strength is soluble in 250 mL or less of aqueous media over the pH range of 1.2 to 6.8, otherwise, low S. Another established classification exists for hepatic extraction ratio (E_H_) - high (>0.7), intermediate (0.3-0.7) or low (<0.3). Except for these, there appears to be no consensus regarding general thresholds for *in vitro* and *in vivo* ADME/PK. Certain threshold have, however, been proposed for *in vitro* microsomal CL_int_ and S (see below) and a P_app_ corresponding to a f_a_ of at least approximately 0.1-0.3 seems generally desired for oral administered drugs. The lack of consensus regarding thresholds for *in vitro* ADME/PK-methods could, at least partly, be explained by differences between methodologies and laboratories and uncertainties regarding *in vitro-in vivo* relationships and what is assumed to be adequate and inadequate.

*In vivo* ADME/PK-profiles could match the ones produced in laboratories. However, appropriate *in vitro* ADME/PK-properties do not necessarily mean that the same will be true *in vivo* in man. For example, a) the combination of low CL_int_ (below, at or near the limit of quantification (LOQ) of hepatocyte and microsome assays) and high P_app_ could imply zero excretion, very slow metabolism and unacceptably long t_½_ *in vivo*, b) the combination of moderate CL_int_ and P_app_ might be associated with low F and short t_½_ *in vivo* if there is high unbound fraction in plasma (f_u_), low steady-state volume of distribution (V_ss_) and considerable excretion, and c) good *in vitro* S could be insufficient for a compound with low permeability and for which high oral exposure and dose are required.

A reliable estimate of oral dose is usually not available in early drug discovery (which makes the interpretation of S-data difficult), and despite efforts at laboratories, a considerable portion of compounds has been shown to have high prediction errors of oral exposure and dose. For example, 16 and 36 % of compounds showed >10-fold prediction errors with the best of selected animal and *in vitro* prediction models, respectively (Poulin et al. 2011), and a near million-fold overprediction of oral exposure and dose has been estimated with classic allometric scaling methodology for one substance (Fuse et al. 2008).

Other obstacles for the screening, selection and prediction process include that ADME/PK for many compounds cannot be quantified at laboratories due to low recovery (Skolnik et al. 2010) and/or low sensitivity (Stringer et al. 2008). The uncertainty for non-quantifiable compounds could potentially imply that they are not considered for further investigation and selection.

For the early drug discovery scientist involved in screening, prediction, ranking and selection of candidate drugs it is important to know how to interpret available ADME/PK-data and decide whether compounds have excellent, sufficiently good or too poor properties and which parameter estimates to improve via design. Based on an apparent lack of consensus regarding thresholds for *in vitro* ADME/PK there is a need for further investigation and clarification.

The objectives of this study were to evaluate variability within and between laboratories for commonly used human *in vitro* ADME/PK methods and to explore whether it is possible to define approximate reliable general thresholds for *in vitro* ADME/PK that could be used for the decision making and compound selection in the early drug discovery process. *In vitro* ADME/PK data for three example drugs (atenolol, diclofenac and gemfibrozil) were used to predict human *in vivo* ADME/PK and investigate whether these would pass a compound selection process.

## Materials & Methods

### *In vitro-in vivo* Data and Relationships

The literature was searched for data for human hepatocyte CL_int_, Caco-2 cell P_app_ and ER, S in water and buffers, intrinsic aqueous S (S_int_; the aqueous solubility of compounds in free acid or free base form, independent on pH and rather more reproducible than other solubility measures) and BCS-class, and for corresponding clinical estimates, including CL_int_ (estimated using the well-stirred liver model and lower blood flow rate of 1500 mL/min), hepatic clearance (CL_H_), E_H_, F, f_u_, f_a_ and maximum *in vivo* dissolution potential (f_diss_; the fraction of oral dose dissolved in the human gastrointestinal tract during optimized conditions). In addition, data on fraction excreted (f_e_) and CYP3A4-specificity (binary classification; substrate/non-substrate) were collected.

Data were taken from the following references - human hepatocyte CL_int_ (Stringer et al. 2008, Bowman and Benet 2019, Hallifax et al. 2010, Sohlenius-Sternbeck et al. 2010, Yamagata et al. 2017, Soars et al. 2002, Cyprotex, 2022), Caco-2 P_app_ (Skolnik et al. 2010; Pham-The et al. 2013, Lin et al. 2011, McGinnity et al. 2007, Bock et al. 2004, Nožinić et al. 2010, Balimane et al. 2006, Hayeshi et al. 2008, Thomas et al. 2005, Matsson et al. 2005, Lee et al. 2017), Caco-2 ER (Skolnik et al. 2010; Pham-The et al. 2013, Lin et al. 2011, McGinnity et al. 2007, Bock et al. 2004, Nožinić et al. 2010, Balimane et al. 2006, Hayeshi et al. 2008), S (Pham-The et al. 2013), S_int_ (Chasing Equilibrium method data (Hewitt et al. 2009, Llinàs and Avdeef 2019)), BCS (Pham-The et al. 2013), CL_H_ and E_H_ (Varma et al. 2010), F (Varma et al. 2010), fa (Pham-The et al. 2013, Varma et al. 2010), fe (Varma et al. 2010), f_u_ (Fagerholm el al. 2021, Wang et al. 2014) and CYP3A4 (Carbon-Mangels and Hutter 2011, Yap and Chen 2005, Bioinformatics 2011, Drugbank). Estimates of f_diss_ were estimated based on f_a_-data (f_diss_≥f_a_) (Pham-The et al. 2013, Varma et al. 2010).

*In vitro-in vivo* relationships, including correlations (R^2^), were established and *in vivo* estimate ranges at certain *in vitro* estimate levels were investigated.

### *In vitro-in vivo* Relationships for Example Compounds

Atenolol, diclofenac and gemfibrozil were chosen as reference compounds for evaluation of *in vitro vs in vivo* ADME/PK-data and thresholds. Atenolol was selected based on moderate *in vivo* permeability and fa (53 %), high S, indications of intestinal efflux, and low *in vivo* CL_int_ (6 mL/min), whereas diclofenac was selected based on its high *in vivo* permeability and f_a_ (99 %) and CL_int_ (58000 mL/min) and uncertainty regarding S. Gemfibrozil was chosen based on observed large interlaboratory variability difference in CL_int_ and f_u_. Differences in results between laboratories have been demonstrated for these three compounds, and especially for gemfibrozil. Their *in vivo* ADME/PK in man was simply predicted considering f_a_ (based on P_app_ and S), CLH (using the well stirred liver flow model with CL_int_ and f_u_), V_ss_ and glomerular filtration rate • f_u_.

### Laboratory Variability

The average laboratory variability for each parameter (both within and between laboratories) was estimated from absolute differences from the mean (*“variability”*) and maximum/minimum-ratios (*“range”*) for the compounds. Contradictory binary classifications were defined as ≤1 *vs* >1.4 for ER, substrate *vs* non-substrate identities for CYP3A4, < or > 10-fold difference for f_u_, and < or > 90 % f_a_ and *in vivo vs in vitro* classing for BCS.

## Results

### *In vitro-in vivo* Data and Relationships

#### Intrinsic Metabolic Clearance

Figure 1 shows the relationships between *in vivo* log CL_int_ (including the median CL_int_ for of 277 drugs = 738 mL/min (Varma et al. 2010)) and E_H_, F and f_e_. There are trends toward increased CL_H_ (R^2^=0.12), E_H_ and CL (R^2^=0.07) with increasing CL_int_, increased F (R^2^=0.04) and f_e_ with decreasing CL_int_, and no apparent relationship between CL_int_ and t_½_ (R^2^=0.009). The median f_e_ at around the median *in vivo* CL_int_ for the drugs (corresponding to an average *in vitro* CL_int_ of ~3 μL/min/10^6^ cells) is ~0.3 (range 0 to 0.8). Near complete renal elimination is also found for many drugs with *in vitro* CL_int_ of ~2 μL/min/10^6^ cells (just above the LOQ of the hepatocyte assay).

**Figure 1.**
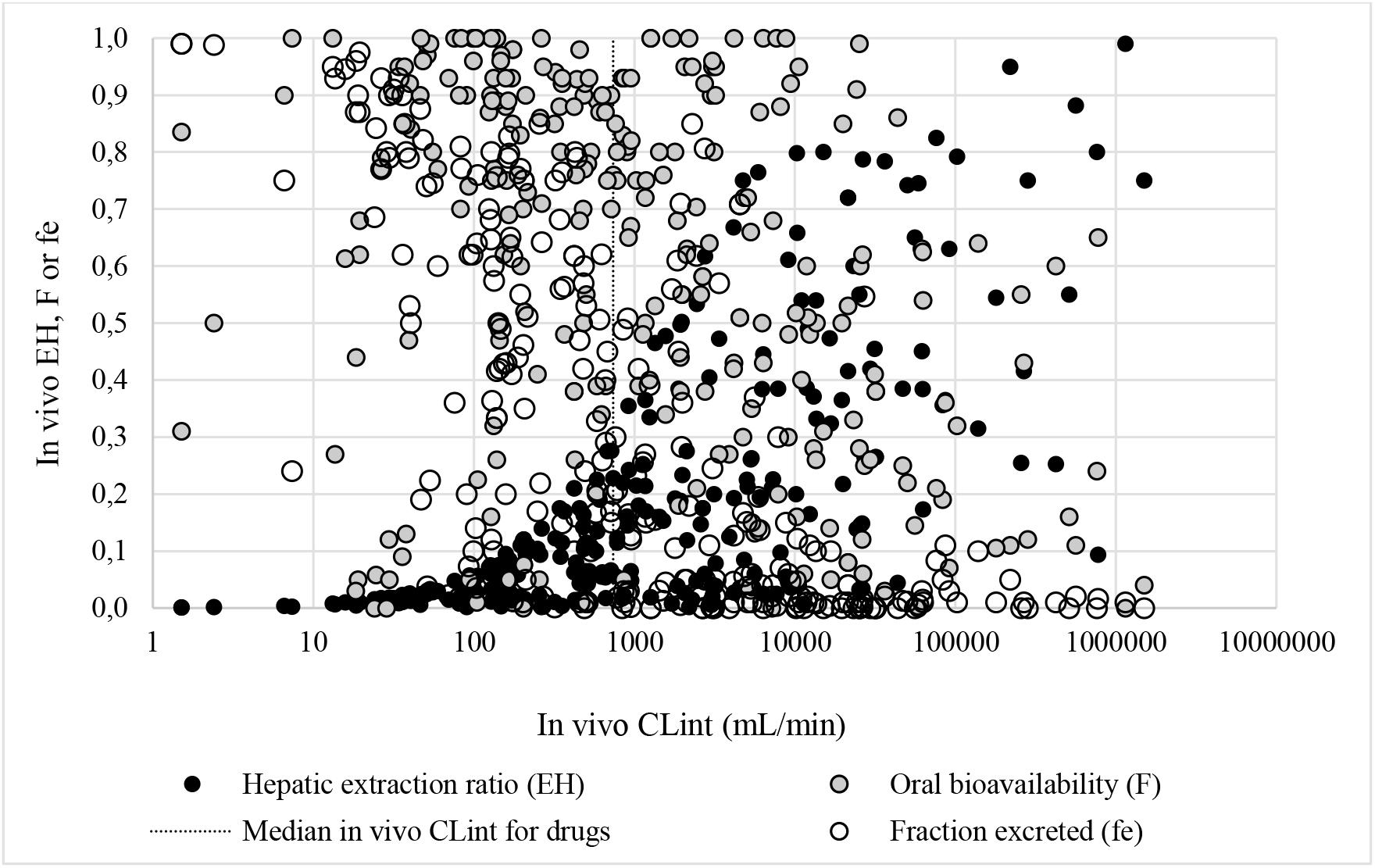
Relationships between *in vivo* log CL_int_ (including the median CL_int_ for of 277 drugs = 738 mL/min and E_H_, F and f_e_.

In Figure 2, the correlation between *in vivo* and *in vitro* log CL_int_ (R^2^=0.44; n=112; (Stringer et al. 2008, Sohlenius-Sternbeck et al. 2010, Yamagata et al. 2017, Soars et al. 2002, Cyprotex, 2022)) is shown. It also shows data for compounds with non-quantifiable CL_int_ (with *in vivo* CL_int_ ranging from 40 to >10000 mL/min). Inclusion of data for 59 compounds with non-quantifiable *in vitro* CL_int_ results in a R^2^ of 0.40. At ~1 (~LOQ) and ~10 μL/min/10^6^ cells, the *in vivo* CL_int_ range ~1500-fold (6.0 to 9136 mL/min) and ~1200-fold (607 to 712500 mL/min), respectively. The *in vivo* CL_int_ for the compounds with quantifiable *in vitro* CL_int_ averages 33000 mL/min and has a median value of 1500 mL/min, which is higher than the median for drugs (738 mL/min). One compound (amiodarone) has a F of 65 % despite a very high CL_int_ (773000 mL/min). Another compound (UCN-01) has a CL of 0.2 mL/min and t_½_ of 1 month (inadequate *in vivo* PK) despite a moderate CL_int_. Low *in vivo* E_H_ is found for 1/3 of compounds with *in vitro* CL_int_≥40 μL/min/10^6^ cells, and high *in vivo* E_H_ is found for 6 % of compounds with *in vitro* CL_int_≤4 μL/min/10^6^ cells.

**Figure 2.**
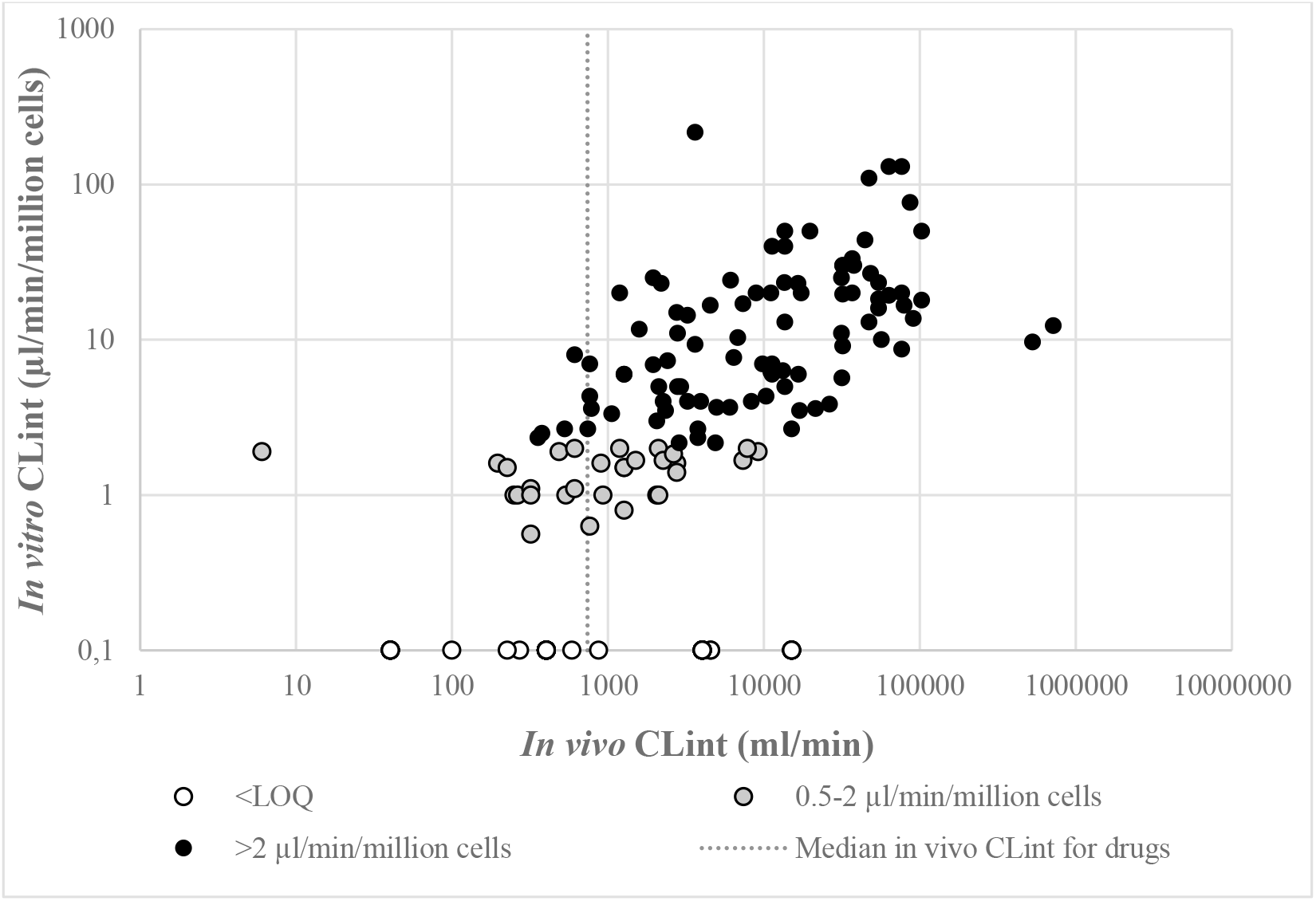
Relationships between *in vivo* and *in vitro* log CL_int_ from various laboratories (R^2^=0.44; n=112; data from 5 sources). The median *in vivo* CL_int_ for of 277 drugs (738 mL/min) is shown. *In vitro* CL_int_ for 59 compounds with non-quantifiable are shown in the figure. Non-quantifiable compounds with *in vivo* CL_int_ <1, 1-10, >10-100 and >100 mL/min/kg from Stringer et al. (2008) were set to have *in vivo* CL_int_ of 40, 400, 4000 and 15000 mL/min, respectively. Inclusion of these non-quantifiable compounds with *in vitro* CL_int_ set to 0.4 μL/min/10^6^ cells resulted in a R^2 of 0.40.

Stringer et al. (2008) investigated intra- and interlaboratory variability for 6 compounds with low to high *in vitro* CL_int_ and found 20 % standard deviation for the former and approximately 2- and 15-fold mean and maximum variability (and average 9-fold range) for the latter. They also found that every other compound had non-quantifiable *in vitro* CL_int_, including ~1/3 of compounds with moderate and high *in vivo* CL_int_. Soars et al. (2002) found a near 4-fold average batch difference and maximum 37-fold range for 8 glucuronidated compounds in their laboratory. Bowman and Benet (2019) compiled hepatocyte CL_int_ data for 50 compounds from 14 papers, and results show ~3-fold mean interlaboratory variability (5-fold mean range) and maximum 51-fold range for one compound (gemfibrozil). Cyprotex showed an overall <2-fold intralaboratory variability (more than half with <1.5-fold difference) for *in vitro* CL_int_ of compounds with *in vivo* CL_int_>2000 mL/min and greater variability for compounds with higher metabolic stability.

#### Apparent Permeability

Pham-The et al. (2013) established a typical sigmoidal relationship between log Caco-2 P_app_ and f_a_ (average of data from various sources; n=282) and found that for Caco-2 P_app_ 1 and 10 • 10^-6^ cm/s the f_a_ averages ~0.6 (range <0.1 to 1) and ~0.8 (range ~0.1-0.2 to 1) and that the standard error is 18.5 %. Several groups/laboratories have found that Caco-2 P_app_ of ~≥10 • 10^-6^ cm/s corresponds to f_a_≥0.8-0.9 (Pham-The et al. 2013). There are also studies which have produced lower (~2-3 • 10^-6^ cm/s (Thomas et al. 2005, Lee et al. 2017)) or higher (~30 • 10^-6^ cm/s (Matsson et al. 2005)) Caco-2 P_app_ at and above this f_a_-limit. A similar trend is apparent for Caco-2 P_app_ corresponding to f_a_=0.5. A Caco-2 P_app_ of ~1-2 • 10^-6^ cm/s often represents a f_a_ of ~0.5, but lower Caco-2 P_app_ is sometimes found at this f_a_-level (~0.1-0.3 • 10^-6^ cm/s (Thomas et al. 2005, Lee et al. 2017)). Weak P_app_-fa-correlations are shown at low Caco-2 P_app_ and low f_a_ (<0.5) (Pham-The et al. 2013, Lin et al. 2011, Matsson et al. 2005).

The approximate average interlaboratory variability for Caco-2 P_app_ is 3-fold (data for 11 compounds in 10 studies (Thomas et al. 2005, Matsson et al. 2005, Lee et al. 2017)). The mean range is 24-fold (Thomas et al. 2005, Matsson et al. 2005, Lee et al. 2017). Corresponding estimates for 5 compounds (atenolol, metoprolol, talinolol, BSP and Gly-Sar) explored at 10 different laboratories were 4- and 44-fold, respectively (Hayeshi et al. 2008). The corresponding intralaboratory values were 2- and 16-fold, respectively (Hayeshi et al. 2008). The standard error for the interlaboratory investigation by Pham-The et al. (2013) was 18.5 %, which is higher than the mean intralaboratory coefficient of variation found by Matsson et al. (2019) (14 %).

Compounds with low recovery (often those with high lipophilicity) seem to be excluded in most/many Caco-2 reports. Sköld et al. (2006) included two such compounds in a Caco-2 P_app_-data set (fenofibrate with log P 5.1 and meclizine with log P 6.2) and reported them as “[*P_app_] not detectable and low recovery*” and “[*P_app_] not reliable due to low recovery*”. Problems to quantify Caco-2 P_app_ and predict f_a_ were also shown by Skolnik et al. (2010), who found that 1/8 and 50 % of compounds were subject to <30 % (44 % of poorly soluble compounds) and <80 % recovery, respectively.

#### Efflux Ratio

There is a trend towards decreased f_a_ with increasing Caco-2 ER (R^2^=0.13; n=207; Figure 3). However, there is no apparent difference in Caco-2 P_app_ *vs* f_a_-relationships for compounds with low ER/no efflux and high ER (Skolnik et al. 2010, Lin et al. 2011). Complete f_a_ (≥0.95) is found in the ER-range 0.28 to 7.0. At ER≥10 the maximum f_a_ is ~0.7, and fa≤0.1 is found for compounds with ER in the range 0.17 to 60.

**Figure 3.**
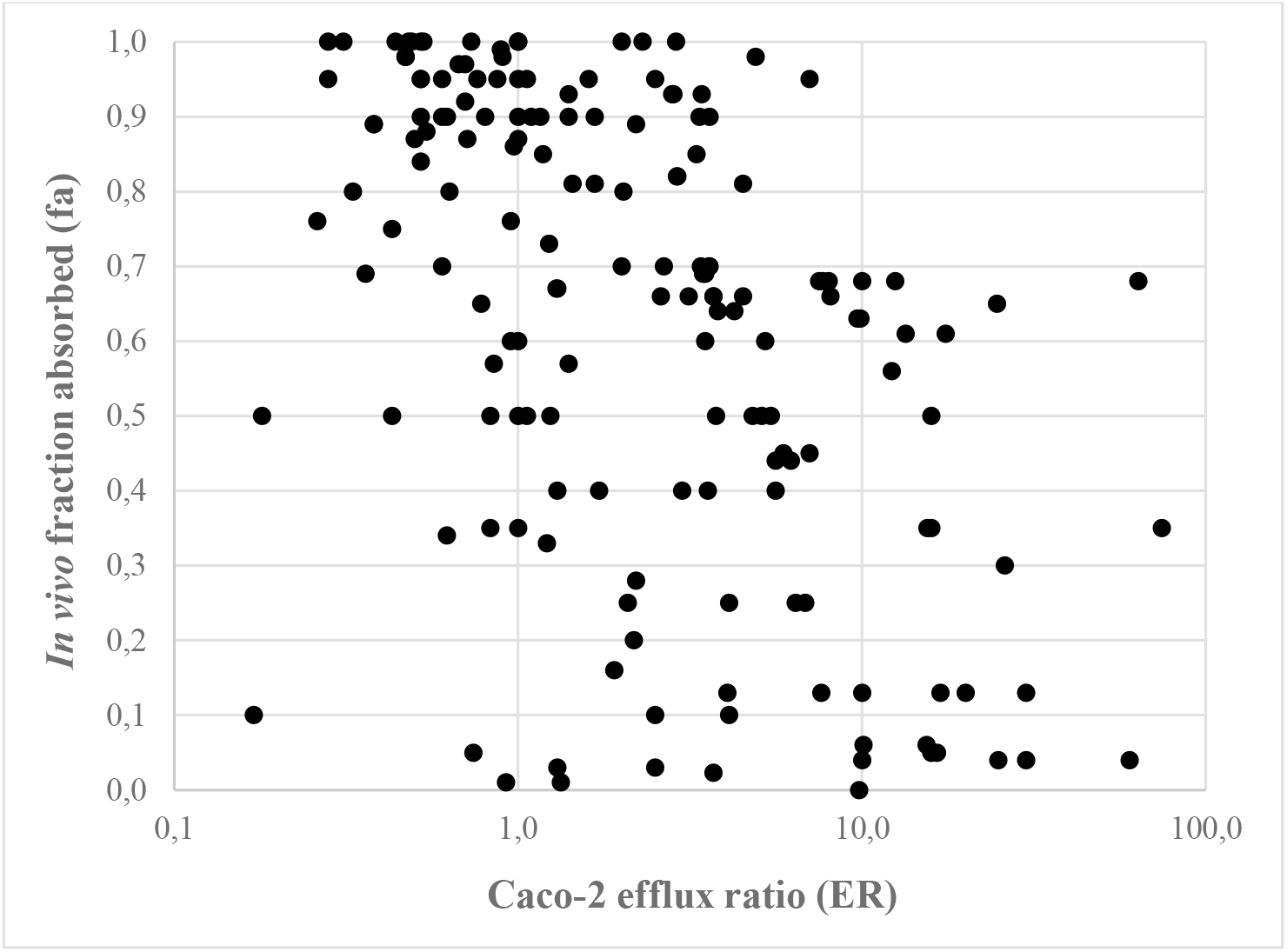
Relationship between Caco-2 cell P_app_ ER from different laboratories and *in vivo* f_a_ (R^2^=0.13; n=207).

Eight of 42 compounds (19 %) have contradictory interlaboratory ER-results (ER≤1 *vs* ER >1.4; substrate *vs* non-substrate for intestinal transporters), and the average, median and maximum range for ER are 5.3- (corresponds to 2.8-fold average variability), 1.7-fold and 93-fold, respectively. Three laboratories out of 9 (33 %) show contradictory ER-results (substrate at one occasion and non-substrate at another) for a compound with moderate Caco-2 P_app_, but only for 1 out of 9 (11 %) for a compound with moderately high Caco-2 P_app_ (Hayeshi et al. 2008). The average intralaboratory range for 5 compounds with mean 1.4 to 20 ER is 1.8-fold (Hayeshi et al. 2008).

#### CYP3A4

CYP3A4-substrate data for were found for 815 compounds, and for 65 (8 %) of these both substrate and non-substrate identities (contradictions) are reported (Carbon-Mangels and Hutter 2011, Yap and Chen 2005, Bioinformatics 2011, Drugbank).

#### Solubility

1, 10 and 100 μM have been suggested minimum S required for minimum acceptable f_a_ of high permeability-compounds at oral doses of 0.1, 1 and 10 mg/kg, respectively (Thomas et al. 2006). Corresponding estimates for low permeability-compounds are ~20, ~200 and ~2000 μM, respectively. A different S-classification was proposed by Hill and Young (2010): >200 μM = high S, 30-200 μM = intermediate S, <30 μM = low S.

S-values collected by Pham-The et al. (2013) shows average, median and maximum interlaboratory range of 100-, 2.3- and 4500-fold, respectively (n=150). The lack of relationship (R^2^=0) between minimum reported *in vitro* S and *in vivo* f_a_ for 40 low S-compounds (S≤0.02 g/L; S ~≤40-80 μM) is shown in Figure 4. Complete or near complete f_a_ is demonstrated for many compounds with S=0.001 g/L (~2-4 μM) and moderate uptake is shown for compounds with S=0.0001 g/L (~0.2-0.4 μM) (Pham-The et al. 2013). Incomplete f_a_ for high permeability compounds appears to be rare. Solubility/dissolution-limited uptake is shown for one highly permeable compound given at high oral doses, griseofulvin (f_a_=45 %; oral dose 250-500 mg).

**Figure 4.**
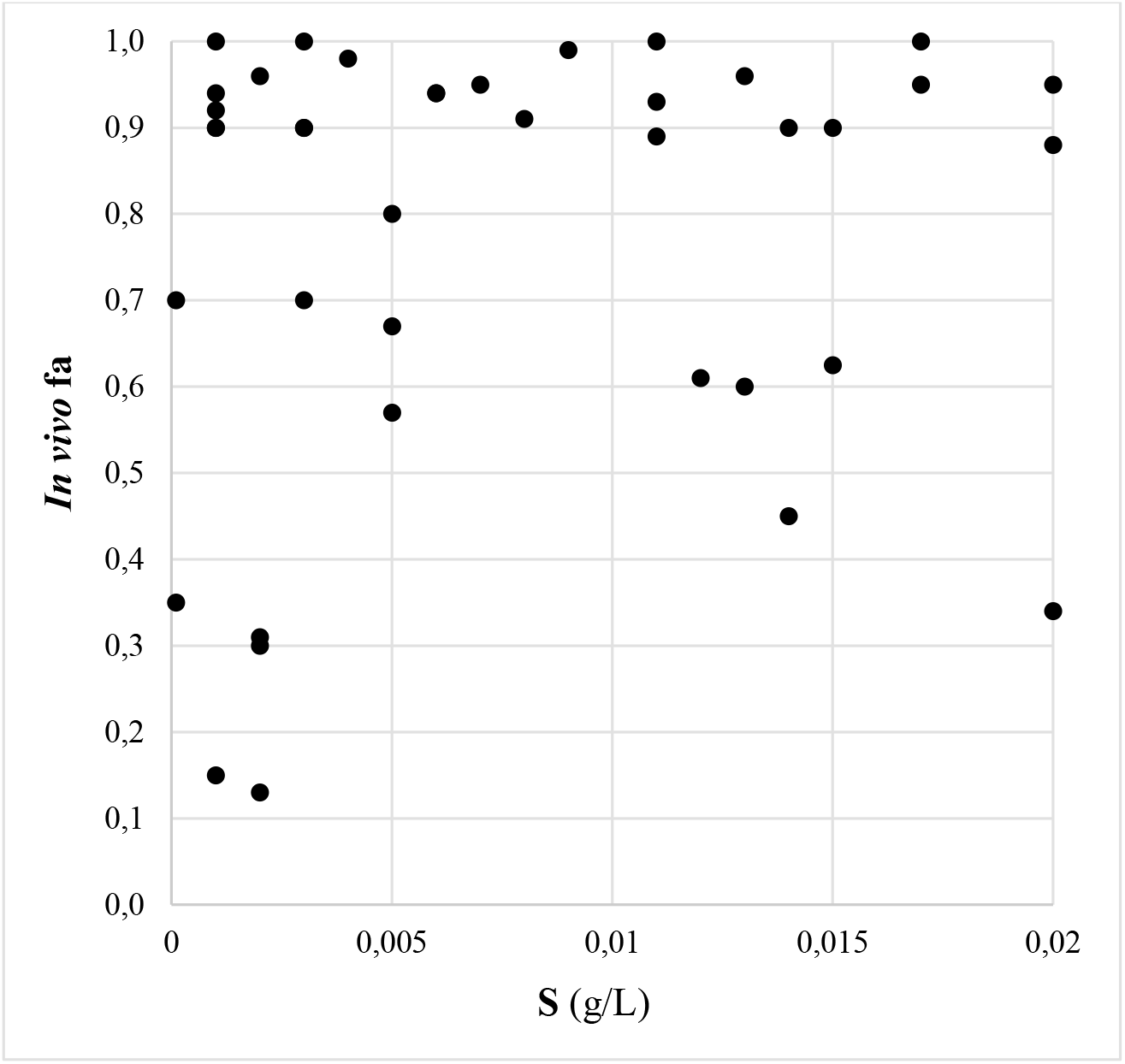
Relationship between minimum reported *in vitro* S and *in vivo* f_a_ for 40 low S-compounds (S≤0.02 g/L; S ca ≤40-80 μM).

Out of 129 compounds with S_int_-data (*Solubility Challenge* data set), 27 have S_int_ in the 1-100 μM-range – amodiaquine, chlorprothixene, diflunisal, dipyramidole, folic acid, glipizide and sulfasalazine in the 1-10 μM-range, and bendroflumethiazide, benzthiazide, carprofen, chlorpromazine, dibucaine, diclofenac, diethylstilbestrol, flufenamic acid, furosemide, imipramine, maprotiline, miconazole, niflumic acid, piroxicam, probenecid, pyrimethamine, sertraline, sulindac, trimipramine and warfarin in the 10-100 μM-range (all of these except folic acid has moderate or high permeability). Only one of these 27 compounds have an *in vivo* gastrointestinal uptake clearly limited/determined by solubility/dissolution (Figure 5).

**Figure 5.**
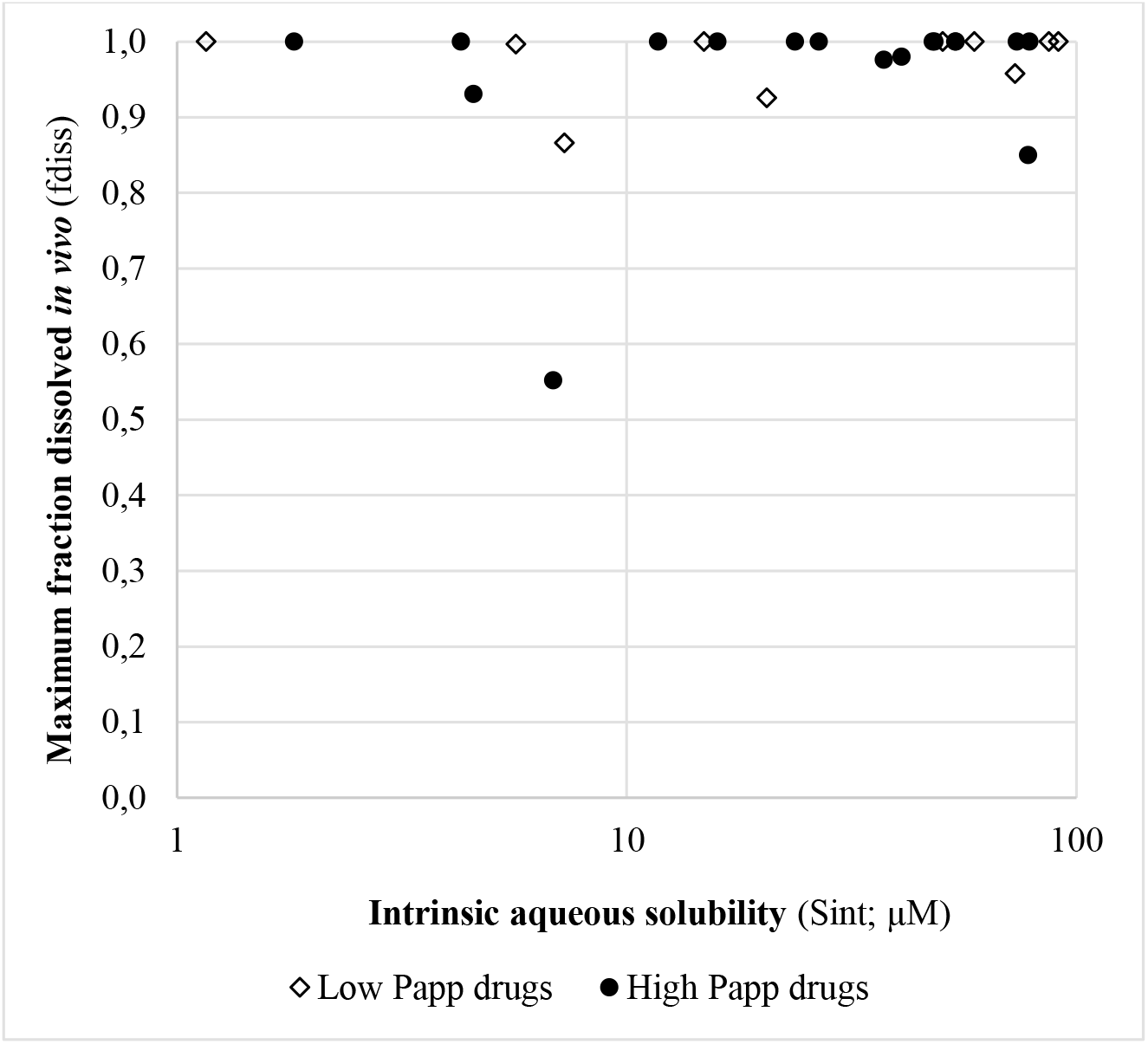
The relationship between *in vitro* log S_int_ and *in vivo* f_diss_ for 27 low S_int_-compounds with high and low P_app_.

The *Solubility Challenge* includes two data sets with S_int_-data for one specific method, one with smaller and one with larger interlaboratory variability (Llinàs and Avdeef 2019) - 1.5-fold standard deviation (corresponding to an approximate average 3-fold variability; n=100) for the former and 4.3-fold standard deviation (corresponding to an approximate average 6.5-fold variability; n=32) for the latter. The maximum individual standard deviation for interlaboratory variability is 8.5-fold (Llinàs and Avdeef 2019).

#### BCS-Classifìcation

Pham-The et al. (2013) collected *in vitro* BCS data for a large set of compounds. For compounds which have BCS-classification from different sources (n=140) 36 % have consistent classification and 64 % show contradictory classes. Most commonly, compounds with undefined *in vitro* BCS-class are classified as class I (high permeability - high solubility) and III (low permeability - high solubility). For compounds with defined BCS-class and predicted/anticipated f_a_ above 90 % (*in vivo* BCS class I) or below 90 % (*in vivo* BCS classes II-IV) (n=73; Figure 5), 37 % are incorrectly classified. 77, 84 and 100 % of compounds predicted/anticipated to be in BCS classes I, III and IV also belong to these classes based on their measured *in vivo* f_a_. For compounds that were predicted/anticipated to belong to BCS class II (high permeability - low solubility), however, only 31 % belong to *in vivo* BCS class II and 69 % have an *in vivo* f_a_ of 0.9-1.0 (BCS class I).

#### Unbound Fraction in Plasma

F_u_-data from various laboratories and with different methodologies were collected for 252 compounds (Figure 7) (Fagerholm et al. 2021). The mean, median and maximum for ratios were 6.8-, 2.0- and 185-fold, respectively (Fagerholm et al. 2021). Corresponding interlaboratory variability estimates were 3.3-, 1.5- and 47-fold, respectively. 5 % of compounds showed >10-fold range for reported f_u_.

**Figure 6.**
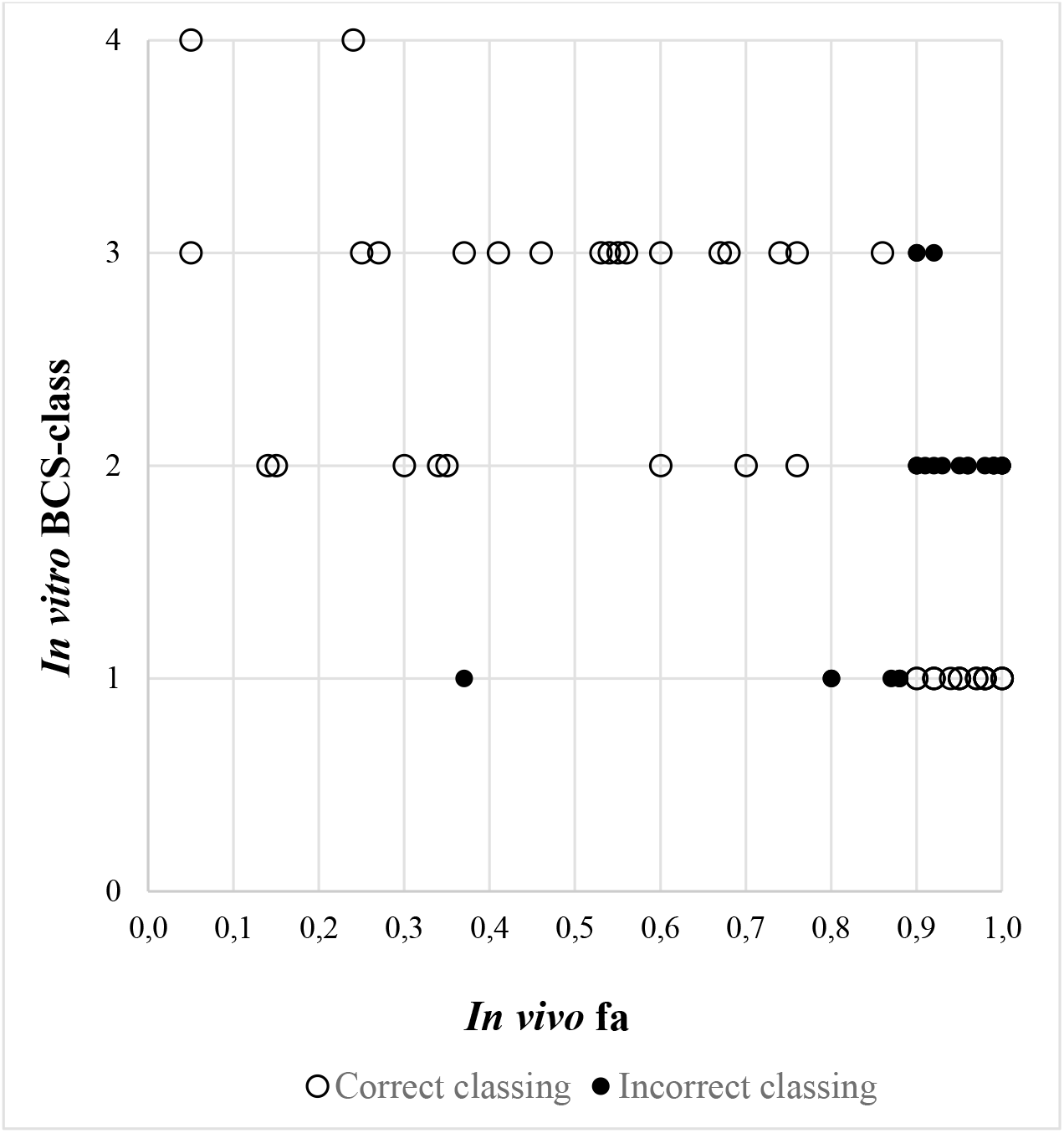
*In vivo* f_a_ *vs in vitro* BCS class for 73 compounds (produced using data from reference 11). The *in vivo* threshold for f_a_ is 0.9 (BCS class I f_a_≦0.9; BCS classes II-IV f_a_≥0.9).

**Figure 7.**
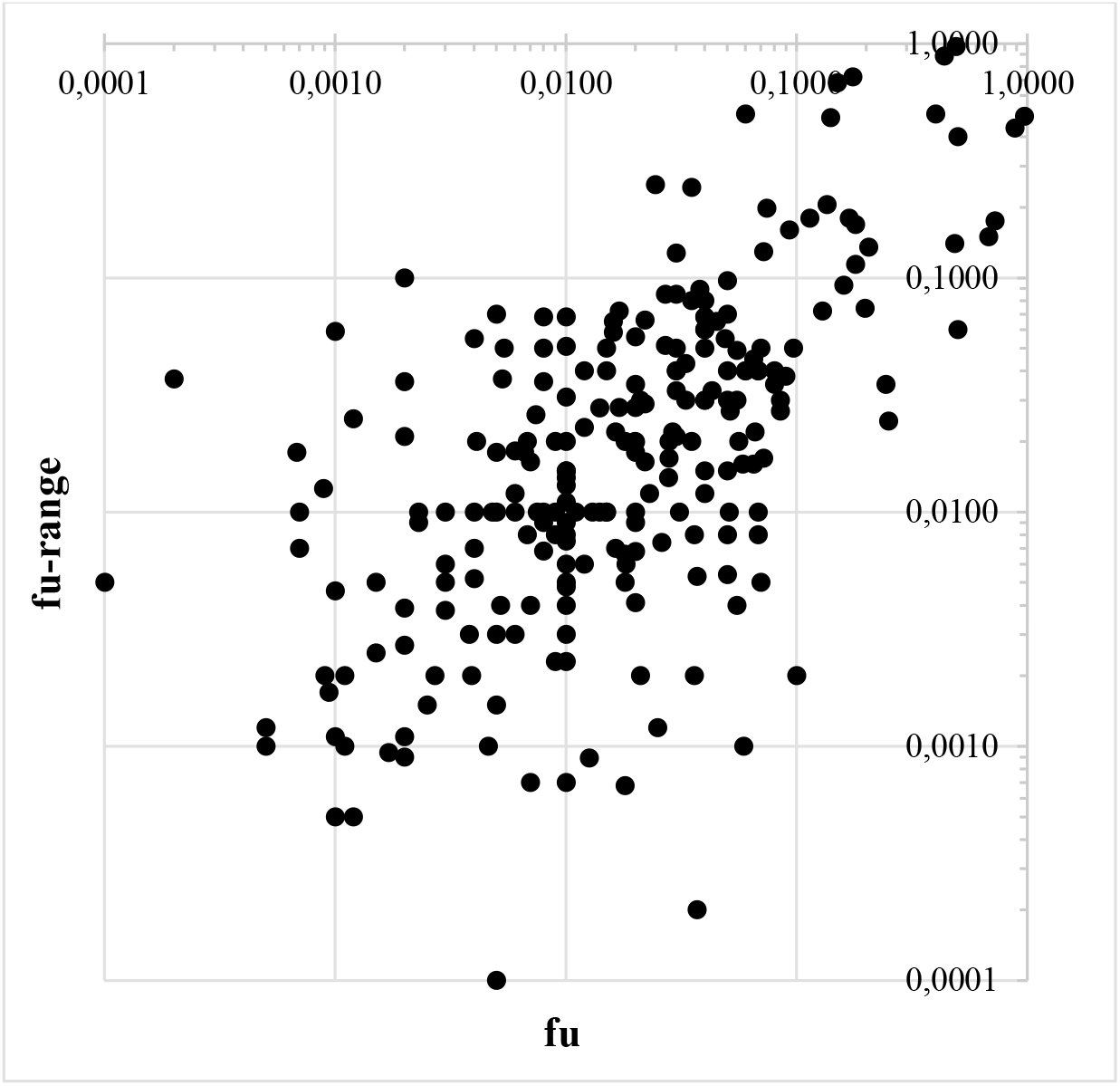
F_u_ *vs* f_u_-range for data collected from various laboratories and with different methodologies (n=252).

Wang et al. evaluated intralaboratory variability for clozapine (mean f_u_=0.13; 4-fold range between laboratories) and warfarin (mean f_u_=0.013; 4-fold range between laboratories) and found ~5 % outliers (maximum f_u_ 0.48 and 0.57 for clozapine and 0.095 and 0.13 for warfarin) (Wang et al. 2014). Intralaboratory variability for clozapine, warfarin, diltiazem, fluconazole, fluoxetine average ~1.1-fold (Wang et al. 2014).

### Predictions for Example Compounds

#### Atenolol

*In vitro* CL_int_ for atenolol has been estimated to 1.9 μL/min/10^6^ cells (Sohlenius-Sternbeck et al. 2010), which corresponds to an average and maximum *in vivo* CL_int_ of ~1700 mL/min and ~18500 mL/min (see Figure 2), respectively. Considering that the *in vivo* CL_int_ is 6 mL/min there is a ~300- fold overprediction of this parameter and an *in vitro* CL_int_ far below the LOQ of the conventional hepatocyte assay is anticipated (Sohlenius-Sternbeck et al. 2010).

Atenolol has a f_a_ of ~50 % (Pham-The et al. 2013). Caco-2 P_app_-values for this compound in the selected reports range between 0.17 (indicating low absorptive capacity) and 11 (indicating high uptake capacity) • 10^-6^ cm/s (65-fold difference), and in 13 of 23 (57 %) cases its reported P_app_ is considerably lower than 1 • 10^-6^ cm/s (McGinnity et al. 2007, Bock et al. 2004, Hayeshi et al. 2008, Thomas et al. 2005, Lee et al. 2017). Largest intralaboratory range shown for atenolol is 16-fold (0.33 and 5.2 • 10^-6^ cm/s; Hayeshi et al. 2008). In one laboratory-comparison study the ER for atenolol averages 1.4-fold and ranges from 0.17 to 3.8 (21-fold difference), with 2/3 of measurements indicating efflux and 4.4-fold maximum intralaboratory difference (Hayeshi et al. 2008). Thus, there is considerable uncertainty (efflux or not; low, moderate or high fa) regarding its predicted gastrointestinal uptake.

The reported f_u_ and S for atenolol are 0.49 to 0.97 and 10000 μM, respectively (Pham-The et al. 2013, Fagerholm et al. 2021). It has moderate F (0.5), determined by moderate uptake capacity (BCS class III - low permeability and high solubility (Pham-The et al. 2013)) and a very low E_H_, intermediate t_½_ (6-7 h), intermediate V_ss_ (0.95 L/kg) and renal excretion as major route of elimination (f_e_=0.99).

According to *in vitro-in vivo* predictions, assuming contribution by glomerular filtration and no tubular reabsorption, intermediate fa, and a V_ss_ of 1 L/kg, atenolol has intermediate E_H_ (~0.5; range ~0.4 to 0.9), low f_e_ (0.13), low/moderate F (~0.25; range ~<0.05 to 0.5) and short t_½_ following intravenous dosing (<1 h; range ~0.5-1.5 h). Absorption-rate determined elimination (“flip-flop”), and longer t_½_ following oral dose administration, is likely.

#### Diclofenac

Reported *in vitro* CL_int_ for diclofenac is high (19 and 125 μL/min/10^6^ cells (Sohlenius-Sternbeck et al. 2010, Cyprotex, 2022), 87 mL/min/kg (Hallifax et al. 2010)), which corresponds to a high *in vivo* CL_int_ (~100000 mL/min). Its reported f_u_ ranges between 0.002 and 0.10 (50-fold interlaboratory variability) (Pham-The et al. 2013, Fagerholm et al. 2021), and its Caco-2 P_app_ is high 13-97 • 10^-6^ cm/s) (McGinnity et al. 2007, Lee et al. 2017). Its reported S ranges from 2 (low) to 9000 (high) mg/L (7-30000 μM) (Pham-The et al. 2013, Llinàs et al. 2008), and the t_½_ is intermediate (1.2-2 h) and V_ss_ is small (0.2 L/kg) (Varma et al. 2010). Despite high CL_int_ and a S_int_ in the risk zone for inadequate gastrointestinal uptake for a highly permeable drug it has a low hepatic CL (260 mL/min), complete f_a_ (0.99) and moderate oral F (0.54).

According to predictions, assuming V_ss_=0.1 L/kg as predicted from animal data (McGinnity et al. 2007) and negligible renal excretion (as for many highly permeable compounds and based on a predicted CL_int_>>glomerular filtration), diclofenac has incomplete (BCS class II if high dose) or complete (BCS class I if low dose) fa, intermediate E_H_ (~0.4; range ~0.1 to 0.9), short t_½_ (0.15 h; range ~<0.1-0.5 h), and moderate F if belonging to BCS class I (~0.6; range ~0.1 to 0.9) or low/moderate F (<0.6) if belonging to BCS class II.

McGinnity et al (2007) used various laboratory data and predicted the t_½_ and daily oral dose of diclofenac in man to be 0.4 h and ~5 grams (~70 times higher than the actual clinical daily dose), respectively.

#### Gemfibrozil

For gemfibrozil, 51- and 18-fold interlaboratory variabilities have been shown for *in vitro* CL_int_ and f_u_ (Bowman & Benet 2019, Fagerholm et al. 2022). The implication of these differences is that two laboratories may show almost 1000-fold differences in CL_int_ • f_u_ (~0.5 *vs* ~450 mL/min). Considering its low E_H_ (for both CL_int_ • f_u_-estimates) and high permeability it is predicted to have high F, high renal tubular reabsorption capacity and minor contribution by renal excretion. Differences in CL_int_ • f_u_ also give almost 1000-fold difference in predicted t_½_. By choosing the low CL_int_ • f_u_ (~0.5 mL/min) there is high potential that the compound would not have passed a stop/go decision.

## Discussion

With the collection of *in vitro* ADME/PK-data from several studies it was possible to make approximations of interlaboratory and intralaboratory variabilities and the extent of non-quantifiable compounds, discrepancies and deviations (Tables 1 and 2). Approximate mean interlaboratory variability for CL_int_, P_app_, ER and f_u_ (3- to 3.5-fold) appears to be about 2- to 3-fold higher than corresponding interlaboratory variability. Mean and maximum interlaboratory range for CL_int_, P_app_, ER, f_u_ and S are approximately 5- to 100-fold and 50- to 4500-fold, respectively, with second largest range for f_u_ and largest range for S. S and f_u_, in particular, are highly sensitive to experimental conditions and presents a serious challenge to laboratories and for human clinical ADME/PK-predictions.

**Table 1.** Approximate variability within and between laboratories for common *in vitro* ADME/PK-methods.

| Parameter | Approximate mean intralaboratory variability (-fold) | Approximate mean interlaboratory variability (-fold) | Approximate mean interlaboratory range (-fold) | Approximate maximum range (-fold) |
| --- | --- | --- | --- | --- |
| Hepatocyte CL<sub>int</sub> | 1.5 | 3 | 5 | 50 |
| Caco-2 P<sub>app</sub> | 2 | 3.5 | 25 | 60 |
| Caco-2 P<sub>app</sub> ER | 1.8 | 3 | 5 | 90 |
| f<sub>u</sub> | 1.1 | 3 | 7 | 185 |
| S | - | - | 100 | 4500 |
| S<sub>int</sub> | - | 3 | - | - |

Intrinsic hepatic metabolic clearance (CL<sub>int</sub>), basolateral to apical/apical to basolateral P<sub>app</sub>-efflux ratio (ER), unbound fraction in plasma (f<sub>u</sub>), apparent permeability (P<sub>app</sub>), solubility (S), intrinsic aqueous solubility (S<sub>int</sub>).

**Table 2.** Approximate non-quantifiable portion and discrepancies within and between laboratories for common *in vitro* ADME/PK-methods.

| Parameter | Approximate non-quantifiable portion (%) (within/between laboratories) | Comment |
| --- | --- | --- |
| Hepatocyte CL<sub>int</sub> | 50 / 50 | <LOQ |
| Caco-2 P<sub>app</sub> | 50 / 50 | <50 % recovery |

| Parameter | Approximate discrepancy (%) (within/between laboratories) | Comment |
| --- | --- | --- |
| Intestinal efflux | 20 / 20 | Caco-2 efflux (Yes/No) |
| CYP3A4-substrate | - / 8 | Substrate ID (Yes/No) |
| BCS-classing *in vitro* | - / 65 |  |
| BCS-classing *in vitro* vs *in vivo* | - / 35 | *In vitro-in vivo* discrepancy |
| f<sub>u</sub> | 5 / 5 | >10-fold difference |

Biopharmaceutics Classification System (BCS), intrinsic hepatic metabolic clearance (CL<sub>int</sub>), unbound fraction in plasma (f<sub>u</sub>), apparent permeability (P<sub>app</sub>).

For a compound with a measured f_u_ of 0.08 in one laboratory other laboratories may (based on results from the collected data) find values of somewhere between ~0.005 and 0.6 for that compound (Figure **7**). Corresponding ranges for compounds with measured f_u_ of 0.01 and 0.5 in one laboratory are ~0.0002 to 0.07 and ~0.05 to 1, respectively. The large variability for f_u_ opens up for retrospective selection of favorable reference f_u_-data for estimation and prediction of, for example, CL_int_, as shown in Fagerholm et al. (2022) where the R^2^ for predicted *vs* observed log CL_int_ varied between <0.1 and 0.8-0.9 depending on choice of f_u_- and CLH-data. The confidence in such predictions may increase by estimating and using mean f_u_-values from various sources (including *in silico* predicted estimates) and comparing to f_u_ (mean and range) for reference compounds.

Variabilities and ranges for measurements of solubility/dissolution are extensive, and these can also be reduced by standardization such as for example using a certain method and protocol (such as S_int_). Despite this, there is questionable applicability of such measurements for predicting and understanding *in vivo* solubility/dissolution. Results by Fagerberg et al. (2015) show a poor correlation between S in phosphate buffer and human intestinal fluid (R^2^<0.2) and, as shown in the present investigation, there is limited/uncertain (actually no) relationship between *in vitro* S and *in vivo* dissolution and uptake (Figures 4 and 5). Proposed minimum S required for achieving minimum acceptable f_a_ (see Thomas et al. 2006) appear exaggerated. A threshold of 1-100 μM for S does not appear adequate. This finding together with the 100-fold average interlaboratory range found for S and comparably high prediction error and uncertainty for predictions of oral exposure and dose implies a major uncertainty/obstacle.

The variability and uncertainty of S-measurements also cause uncertainty and contradictory results for BCS-classification. 2/3 and 1/3 discrepancies between studies for *in vitro* BCS-classification and *in vitro* and *in vivo* BCS (Figure 6), respectively, are remarkable. Most compounds predicted to belong to BCS class II belong to *in vivo* BCS class I, which is in line with exaggerated solubility limitations as shown and discussed above and in reference 36.

Interlaboratory variability is also quite pronounced for the Caco-2 assay, both for P_app_- and ER- measurements. The variability and uncertainty were demonstrated for atenolol. Based on P_app_-measurements it is not clear whether this compound is effluxed or not or is predicted to have low, moderate or high f_a_. In many cases, Caco-2 P_app_ of ~1-2 and ≥10 • 10^-6^ cm/s represent f_a_ of ~0.5 and ~≥0.8-0.9, respectively. Overall, a Caco-2 P_app_ of ~≥2-3 • 10^-6^ cm/s seems compatible with high level of permeability-determined gastrointestinal uptake (f_a_≥0.8-0.9). The use of P_app_ for reference compounds with known f_a_ is recommended for calibration/adjustment for the interlaboratory variability along the P_app_-scale (up to ~±3- to 10-fold) of P_app_ *vs* f_a_-relationships. Otherwise it is difficult to tell whether a P_app_ in the approximate range 0.1 to 2 • 10^-6^ cm/s is sufficiently high for assuring/predicting adequate gastrointestinal uptake. At lower P_app_ it is even more difficult to predict whether a compound will be absorbed to more than, for example, 10-30 %.

Efflux influences the extent of gastrointestinal uptake, at least and in particular for compounds with low and moderate P_app_. An ER>7-10 seems to be associated with an incomplete f_a_ (Figure 3), but besides that the magnitude of ER appears to have limited impact on f_a_-predictions. Sufficiently high f_a_ can be reached despite an ER exceeding 70. This and a ~90-fold maximum interlaboratory range makes it difficult to set a useful and clear threshold for ER. 20 % discrepancies (both efflux and non-efflux shown for a compound) within and between laboratories imply additional uncertainties.

Interlaboratory discrepancy is also demonstrated for CYP3A4-substrate identity, and it appears to be the lowest of those studied and compared in this study (8 %).

Low recovery is another important and common (and generally not often covered and discussed) issue with the Caco-2 assay. In such cases (apparently, for almost every other compound), *in silico* methodology may be particularly useful for predictions of f_a_.

The variability, range and coverage for the hepatocyte assay appear to be of the same size as for Caco-2 – 1.5- to 2-fold intralaboratory variability, 3- to 3.5-fold interlaboratory variability, 50- to 60-fold maximum range and approximately 50 % quantifiable compounds. Improvements can be achieved by the inclusion of reference compound data and use of *in silico* methodology (with higher sensitivity and capacity to predict *in vivo* CL_int_ of compounds with *in vitro* CL_int_<LOQ) and a hepatocyte assay capable of quantifying low CL_int_-compounds.

Thresholds for *in vitro* CL_int_ have been proposed and used (20 and 25 μL/min mg protein for microsomes; corresponds to an approximate moderately high average *in vivo* CL_int_ of 7000 mL/min (Lee et al. 2007, Sakiyama et al. 2008, Liu et al. 2015, Hu et al. 2010).To set sharp thresholds for *in vitro* CL_int_-values seems, however, difficult, especially such that are clinically relevant and useful. Reasons include the facts that a) the *in vitro* LOQ represents low up to moderately high *in vivo* CL_int_ (up to >10000 mL/min; >7-fold the liver blood flow rate), b) there is several thousand-fold variability of *in vivo* CL_int_ for compounds with an *in vitro* CL_int_ of ~1 μL/min/10^6^ cells (~LOQ), c) hundred- to thousand-fold predictions errors (overpredictions of *in vivo* CL_int_; as for atenolol) have been produced with the hepatocyte assay (Sohlenius-Sternbeck et al. 2010), d) a low CL_int_ is not necessarily favorable (risk of very long t_½_ for a compound with low f_u_ and high P_app_), e) there are poor correlations (R^2^=~0-0.1) between *in vivo* CL_int_ and CL_H_, CL, t_½_ and F, f) sufficiently high F and long t_½_ are achievable despite very high CL_int_ (including compounds with *in vivo* CL_int_ of nearly a million mL/min and ~100000 mL/min (diclofenac)), and g) inadequate human clinical PK has been found despite moderate-size CL_int_ (as for UCN-01). The correlations between *in vivo* CL_int_ and CL_H_ (R^2^=0.12) and CL (R^2^=0.07) are weak. CL_int_ needs to be considered together with other parameters such as f_u_, extrahepatic elimination (often significant for compounds with *in vitro* CL_int_ of same size as LOQ) and Vss, and inclusion of *in vitro* CL_int_ for reference compounds with known *in vivo* CL_int_ is recommended.

The findings challenge *in silico* methods based on *in vitro* ADME/PK-data, for example, for S, f_a_ and binary CL_int_. It is possible to reach 70-80 % correct binary *in silico* predictions of *in vitro* microsome CL_int_ using a threshold of 20-25 μL/min mg protein (corresponds to an *in vivo* CL_int_ ranging from ~500 to 25000 mL/min (Lee et al. 2007, Sakiyama et al. 2008, Liu et al. 2015, Hu et al. 201))), but the clinical relevance of such models is not obvious. *In silico* models for S are hindered by the lack of apparent correlation between *in vitro* S and *in vivo* f_diss_.

The predictions of human clinical ADME/PK for the selected reference compounds showed that atenolol and diclofenac (and dependent on choice of data, maybe also gemfibrozil) lacked sufficiently good/promising properties and would be at risk of being stopped from further development (in case they were candidate drugs). *In vitro* ADME/PK-data for atenolol and diclofenac are varying and include either poor permeability, solubility and/or metabolic stability. Integrated predictions showed low F and short t_½_. The outcomes show the importance to consider laboratory variability, include reference compounds and uncertainty measurements, validate prediction methods, and consider extrahepatic elimination (including gut-wall degradation and renal and biliary excretion) and absorption rate in predictions, and that one should be careful in basing choices and making decisions on thresholds.

Limitations with the study include that it is not based on all reports with useful data (it is based on data from a selection of important papers in the field), that approximations (rather than exact estimations) of variability were made and ADME/PK-predictions for atenolol, diclofenac and gemfibrozil were incomplete and relatively simple (as usually during the early discovery phase). Despite this, results show the variabilities and uncertainties with laboratory ADME/PK-data. Addition of data may not dramatically change the overall picture presented here.

## Conclusion

The overall impression for *in vitro* ADME/PK methods is that interlaboratory variability is considerable, especially for f_u_, S, ER and BCS-classification, and on average about twice as high as corresponding variability within laboratories, and that it appears difficult or impossible to set clear clinically useful thresholds, especially for CL_int_, ER and S. Caco-2 P_app_ appears to be most suitable for setting a threshold for sufficient and good properties. Most notable is the very large variability and poor/absent *in vivo* correlation for S, poor *in vitro-in vivo* consistency for BCS-classification, large portion of compounds out of reach for Caco-2 and hepatocytes, and predictions for atenolol, diclofenac and gemfibrozil suggesting that they would/might have inadequate ADME/PK-characteristics *in vivo* in man (and therefore opted out as suitable candidate drugs). Improvements are believed to be achievable with the inclusion of average data for reference compounds, use of prediction methods for extrahepatic elimination and complementary *in silico* prediction methodology with wider range and higher accuracy, and an increased insight into the relationship between *in vitro* S and *in vivo* solubility/dissolution.

